## Supplementary Figures 1 - 10 for "PTMNavigator: Interactive Visualization of Differentially Regulated Post-Translational Modifications in Cellular Signaling Pathways"

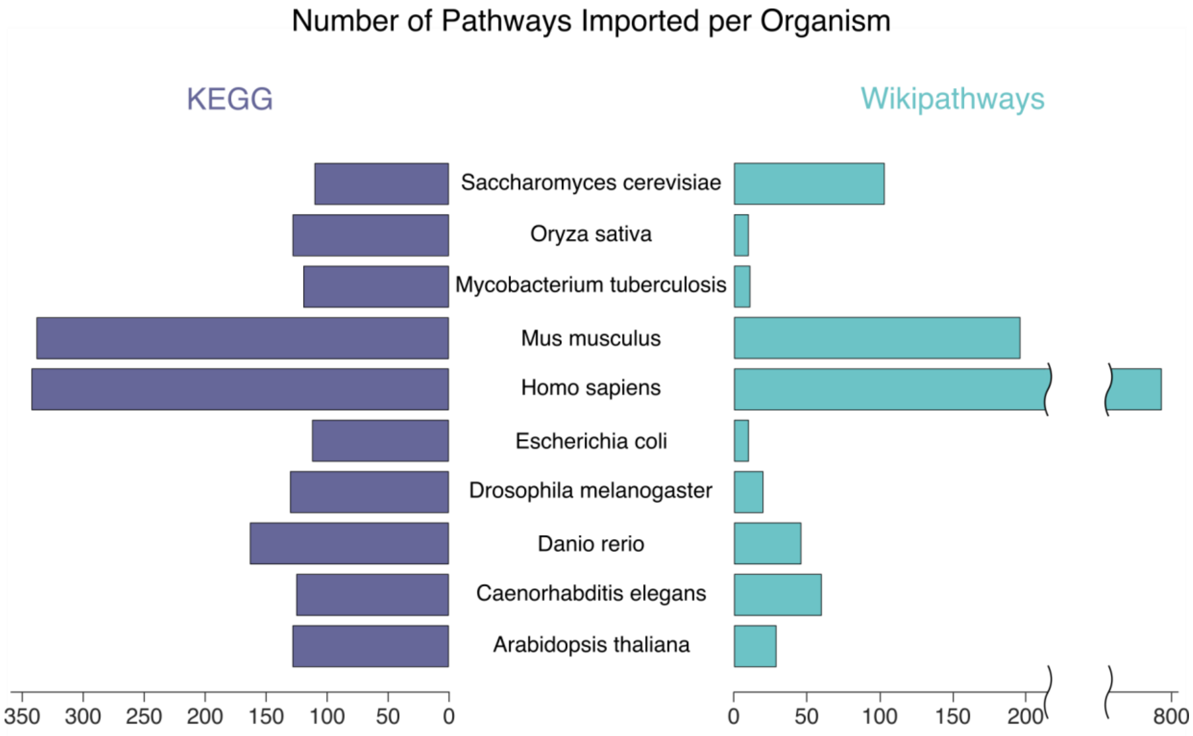


**Fig. S1. Number of pathways per organism and reference database that were imported into ProteomicsDB and are available in PTMNavigator.**

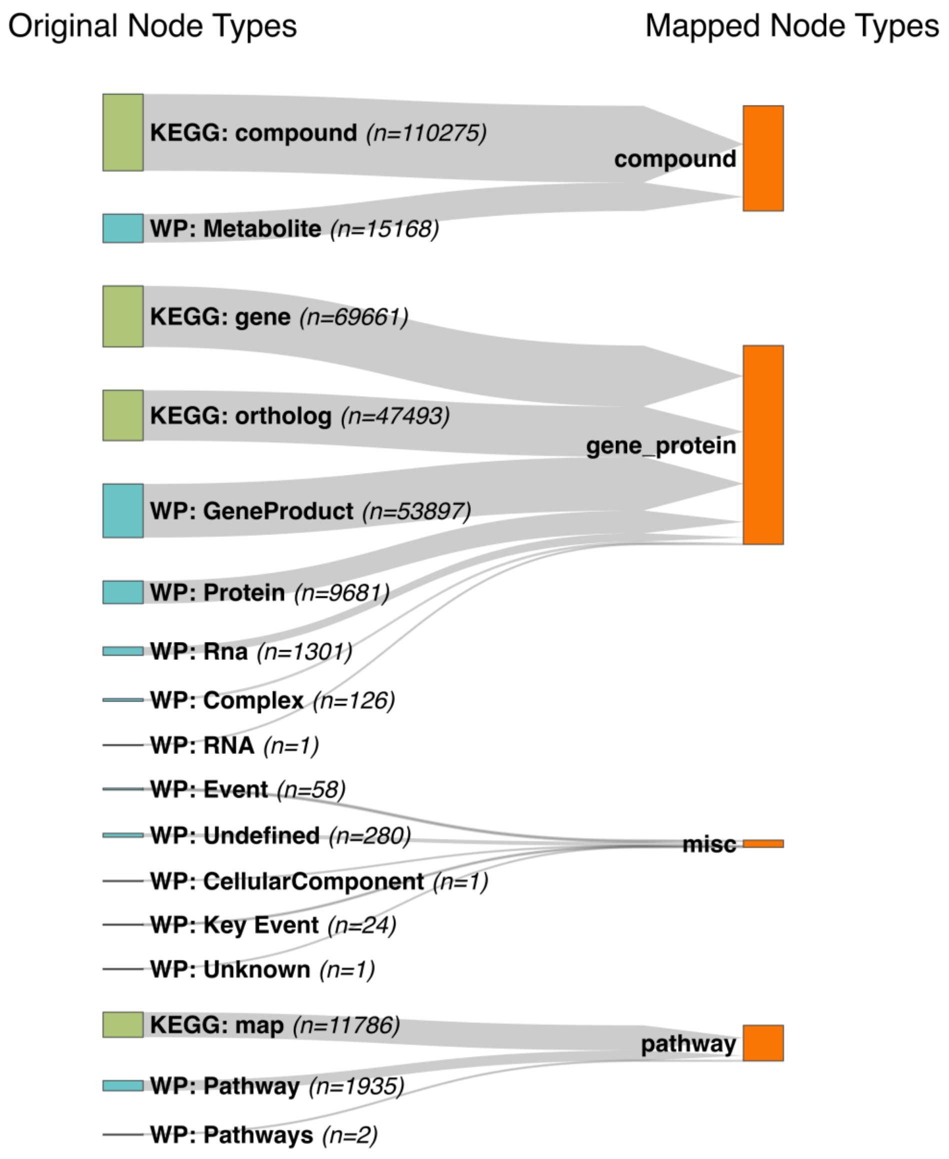


**Fig. S2. Mapping of Node Types from KEGG and WikiPathways to PTMNavigator.** On the left side all node types are listed that appear in the KEGG KGML or WikiPathways GPML files of the ten imported organisms. On the right side are the 4 node types that are defined in PTMNavigator.

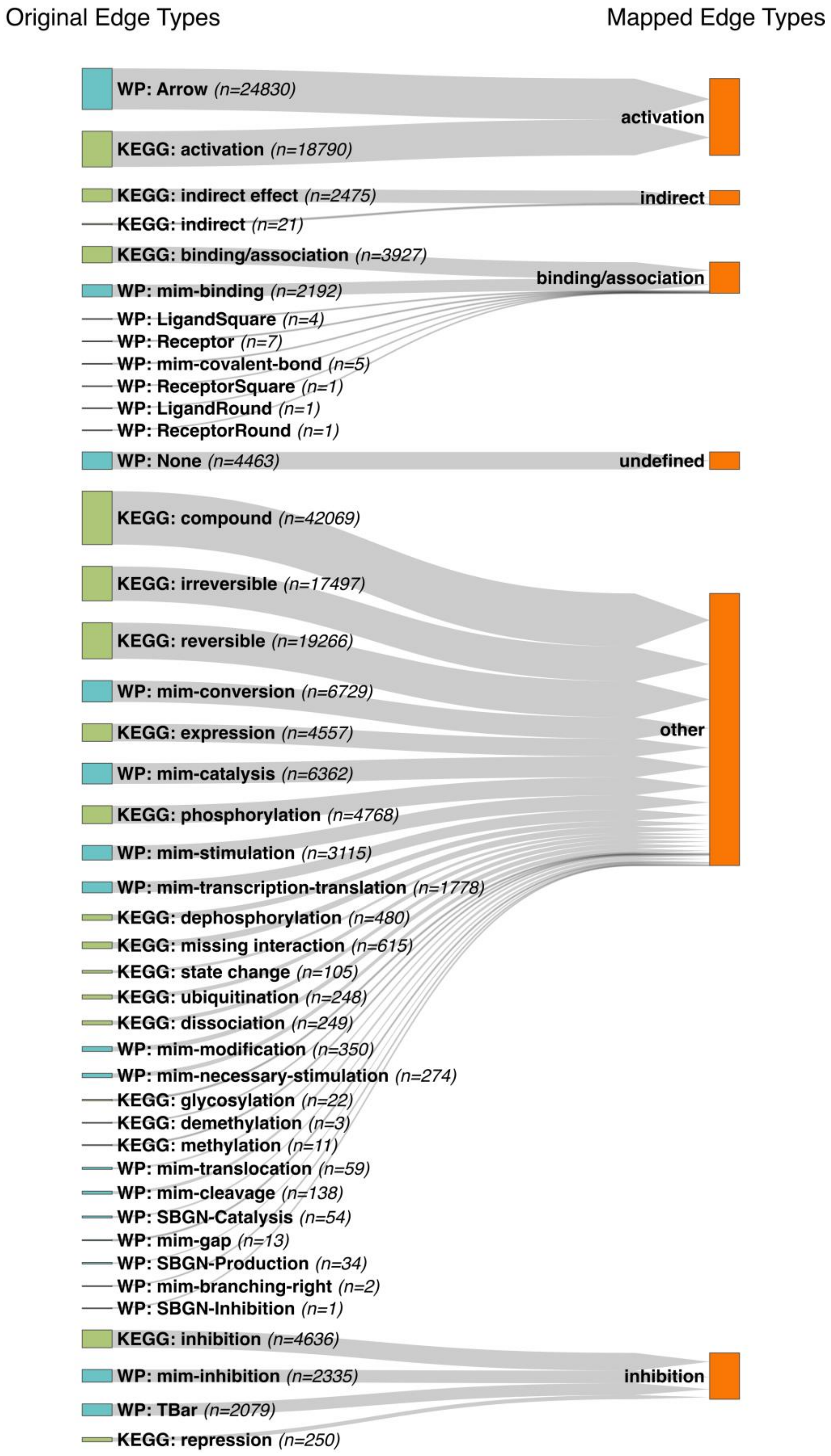


**Fig. S3. Mapping of Edge Types from KEGG and WikiPathways to PTMNavigator.** On the left side all edge types are listed that appear in the KEGG KGML or WikiPathways GPML files of the ten imported organisms. The right side shows how they are mapped to the internal JSON representation in ProteomicsDB. Raw edge types are converted either to ‘activation’, ‘inhibition’, ‘binding/association’, ‘indirect’ or ‘undefined’. The other edges are not converted and retain their original type (in the figure, these edge types are summarized as ‘other’).

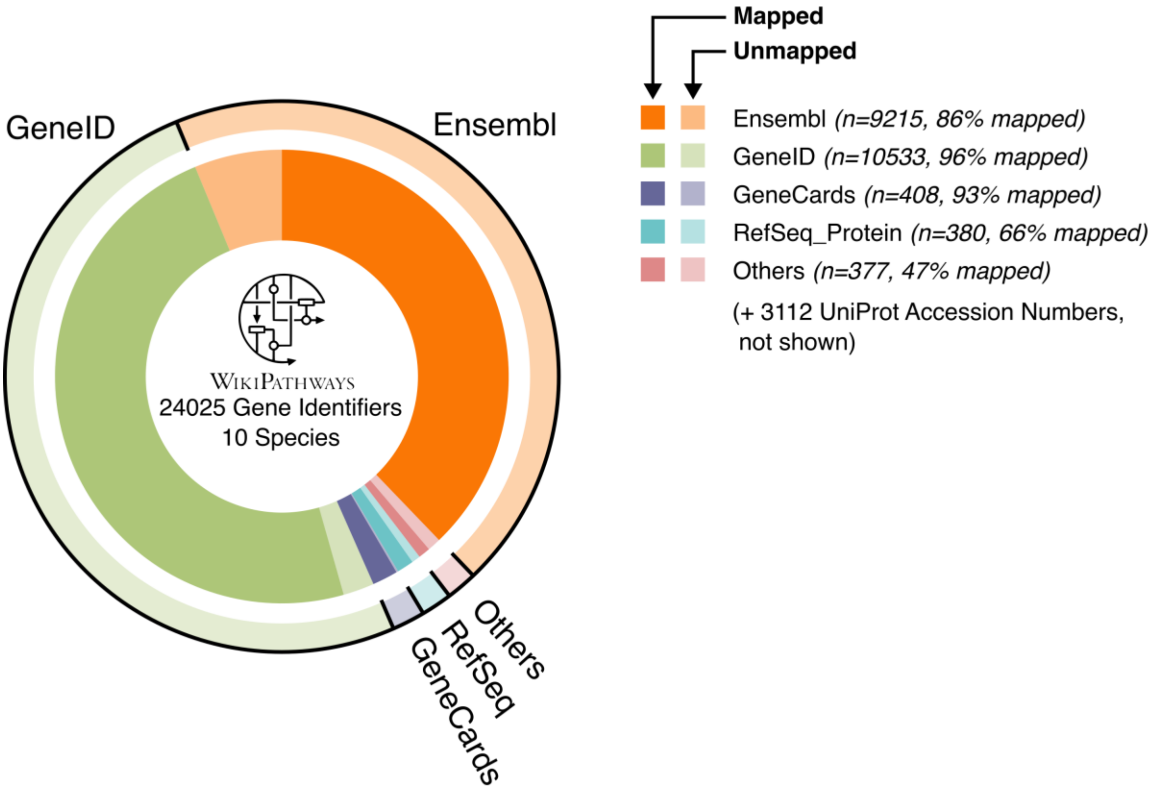


**Fig. S4. Gene Identifier Conversion for WikiPathways Diagrams.** WikiPathways uses 9 different gene identifier types in the diagrams processed in this study. The UniProt API was used to map them to UniProt Accession Numbers. Overall, 91% of gene identifiers (21956 of 24025) could be mapped (or were already in UniProt format). 'Ensembl' summarizes identifiers of type 'Ensembl' and 'Ensembl_Transcript'. 'Others' summarizes 'KEGG', 'EC', and 'HGNC' identifiers.

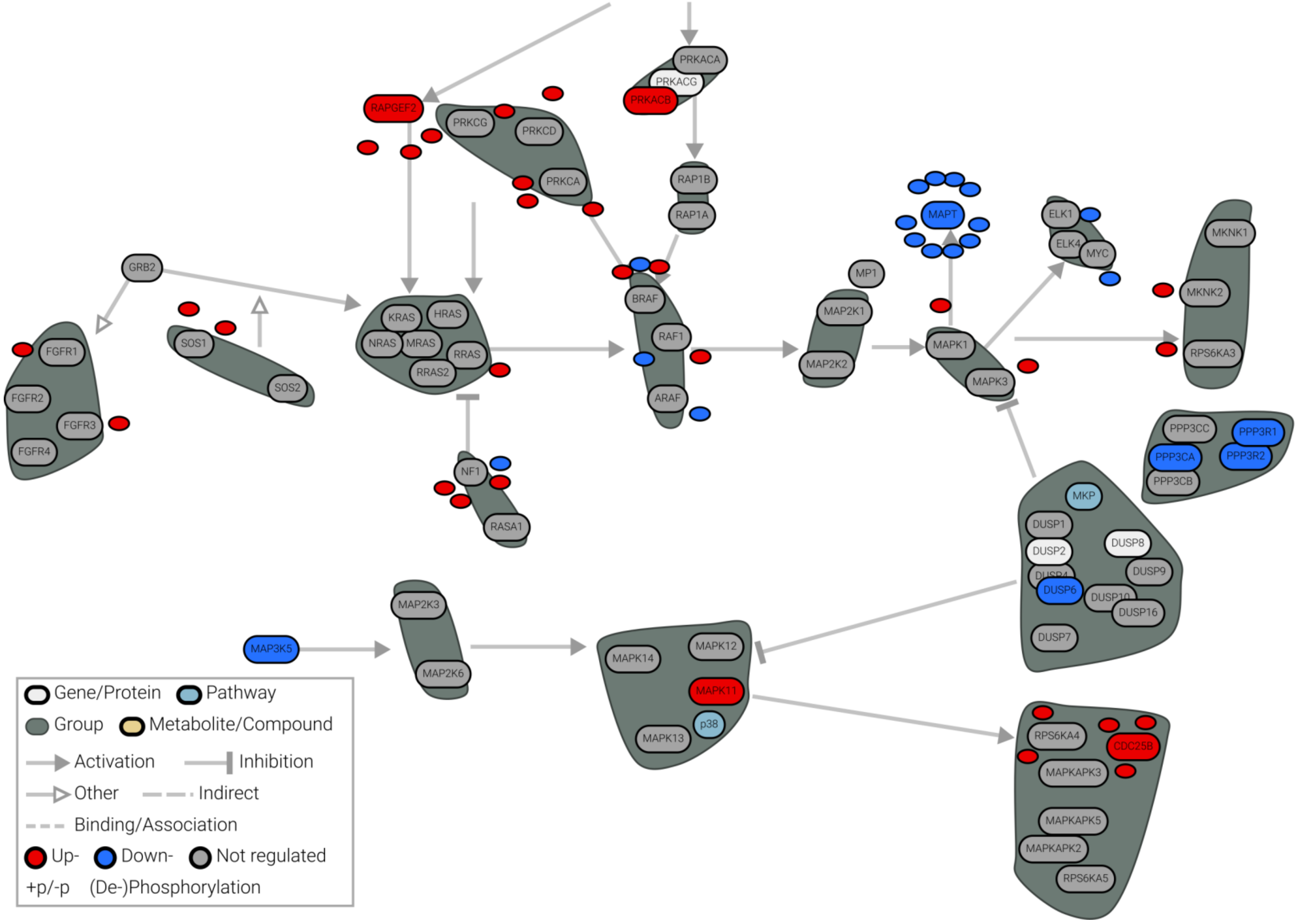
RAPGEF2
PRKCG
PRKCD
PRKCA
PRKACA
PRKACG
PRKACB
RAP1B
RAP1A
GRB2
SOS1
SOS2
FGFR1
FGFR2
FGFR3
FGFR4
KRAS
HRAS
NRAS
MRAS
RRAS
RRAS2
NF1
RASA1
BRAF
RAF1
ARAF
MP1
MAP2K1
MAP2K2
MAPT
MAPK1
MAPK3
ELK1
ELK4
MYC
MKNK1
MKNK2
RPS6KA3
PPP3CC
PPP3R1
PPP3CA
PPP3R2
PPP3CB
MKP
DUSP1
DUSP2
DUSP8
DUSP9
DUSP6
DUSP10
DUSP16
DUSP7
MAP3K5
MAP2K3
MAP2K6
MAPK14
MAPK12
MAPK11
p38
MAPK13
RPS6KA4
CDC25B
MAPKAPK3
MAPKAPK5
MAPKAPK2
RPS6KA5
Gene/Protein
Pathway
Group
Metabolite/Compound
Activation
Inhibition
Other
Indirect
Binding/Association
Up-
Down-
Not regulated
+p/-p
(De-)Phosphorylation

**Fig. S5: Visualization of protein-level and peptide-level data in PTMNavigator.** Comparison of baseline proteomes and phosphoproteomes of the cell lines A204 and RDES projected onto the diagram 'MAPK Signaling - WP382'. Colors indicate whether expression is higher in A204 (red) or RDES (blue). Grey color indicates peptides/proteins that were not sufficiently different between the two cell lines (as thresholds, a difference of 3 was used for proteomic and 4 for phosphoproteomic data). White protein nodes indicate proteins that were not measured in the experiment. Dataset generated by Lee et al.

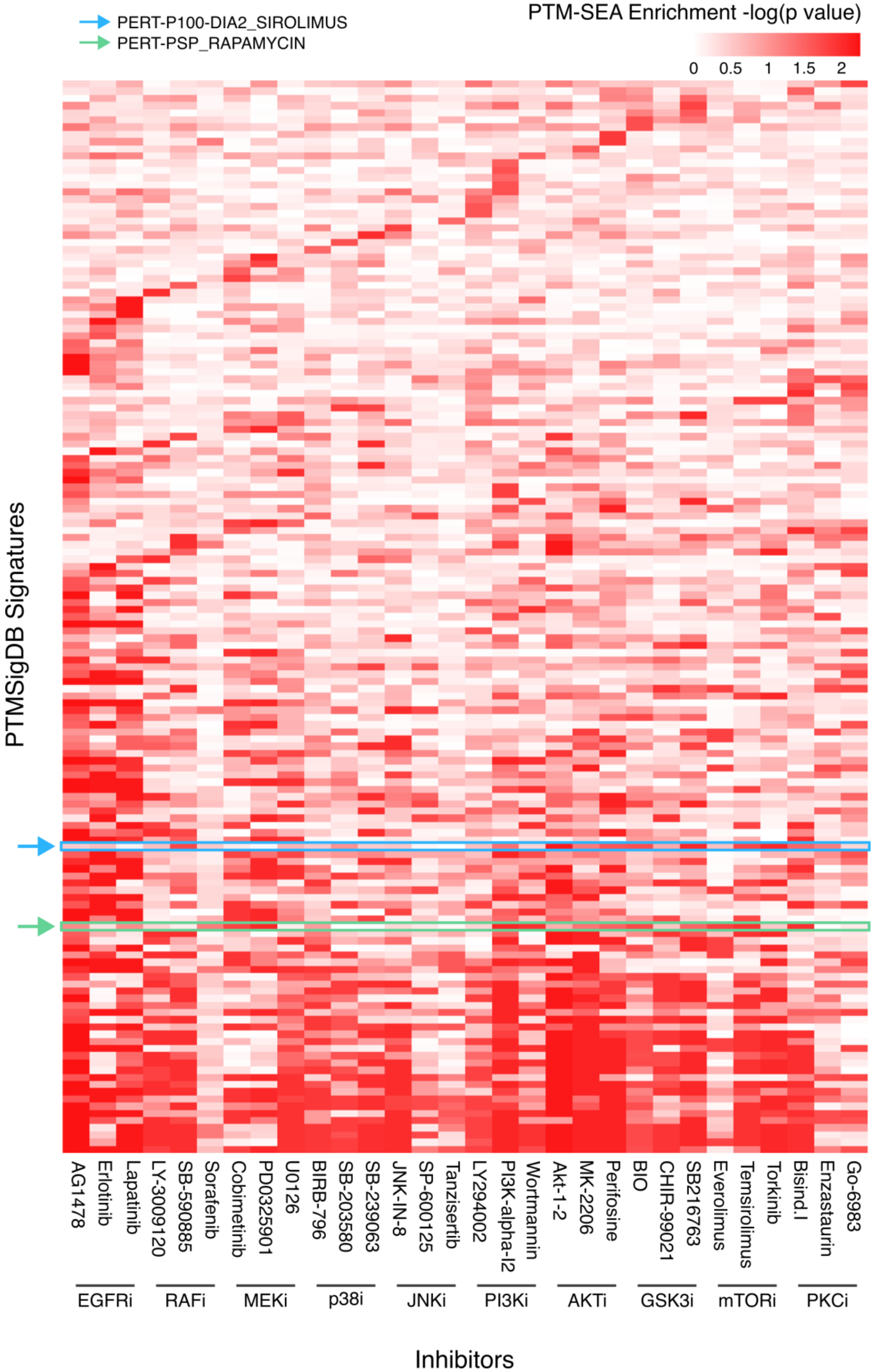


**Fig. S6. Results of PTM-SEA analysis of the Bekker-Jensen dataset.** Shown are signatures with p-value<0.05, sorted by the number of datasets in which they were reported as enriched (larger numbers on the bottom). For illustrative purposes, only signatures that are significant for at least 6 datasets are shown. Inhibitors are grouped by their designated targets.

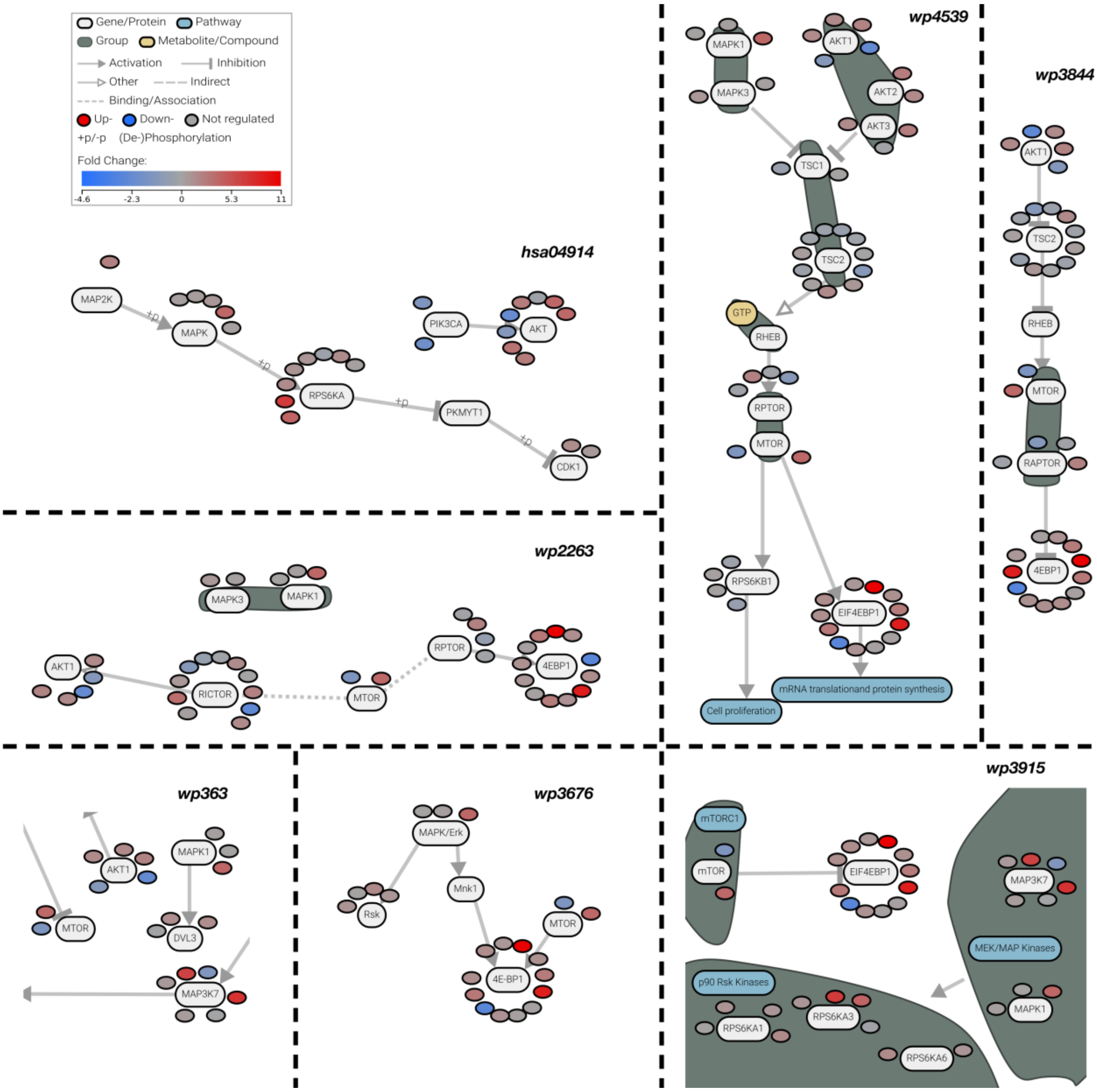


**Fig. S7. Common subnetworks of the high-scoring pathways in the Go-6983 dataset.** Edited Screenshots from PTMNavigator showing excerpts from 7 different canonical pathway diagrams. All of these pathways showed significant enrichment in the gene-centric-redundant ssGSEA. PTM nodes are colored according to their fold change. Edges labeled with '+p' indicate a phosphorylation of the target protein.

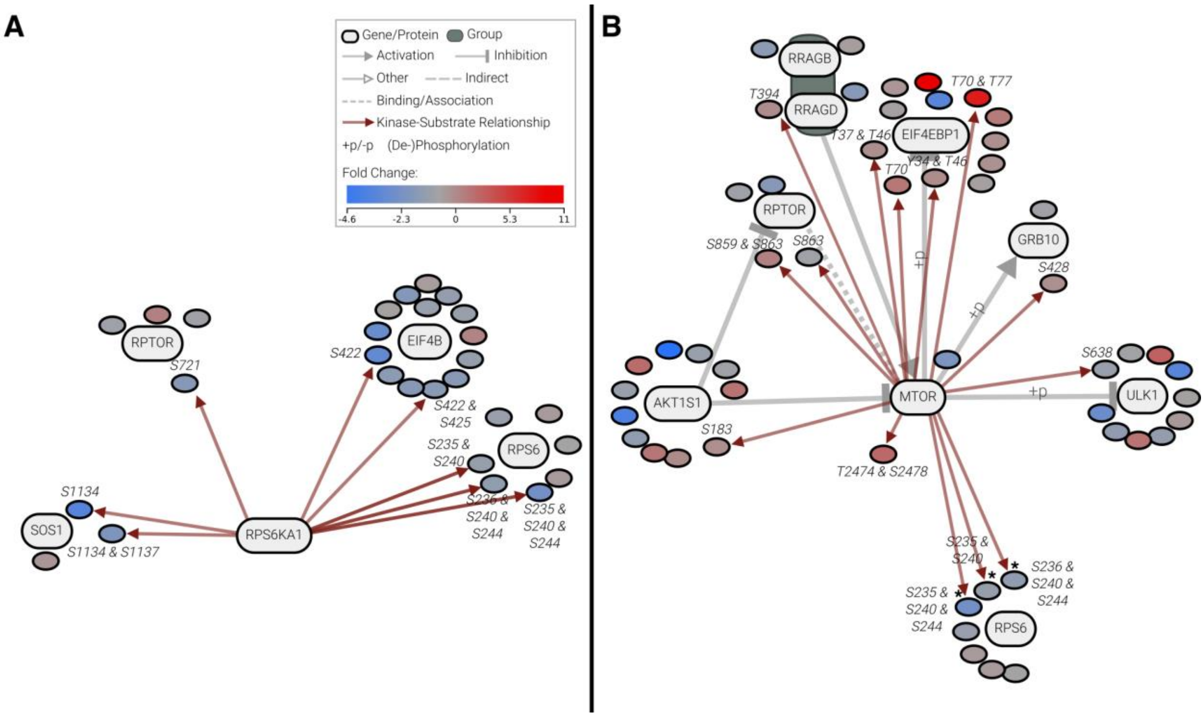


**Fig. S8. PTMNavigator makes PTM-SEA and kinase activity scores transparent.** Edited Screenshots from PTMNavigator showing excerpts from the projection of the Go-6983 dataset onto the pathway 'mTOR Signaling Pathway - hsa04150'. Shown are the proteins phosphorylated by RPS6KA1 (**A**) and mTOR (**B**). Red arrows indicate kinase-substrate annotations from PhosphoSitePlus. Both PTM-SEA and KSEA reported a downregulation of RPS6KA1 activity as well as an upregulation of mTOR activity. The visualizations of PTMNavigator confirm this, with all annotated RPS6KA1 substrates in the diagram showing reduced expression, whereas most mTOR substrates showed increased expression. Notably, the sites on RPS6 (indicated by asterisks '*') are not exclusively associated with mTOR, but are also substrates of RPS6KA1, RPS6KA3, and PRKCD. Edges labeled with '+p' indicate a phosphorylation of the target protein.

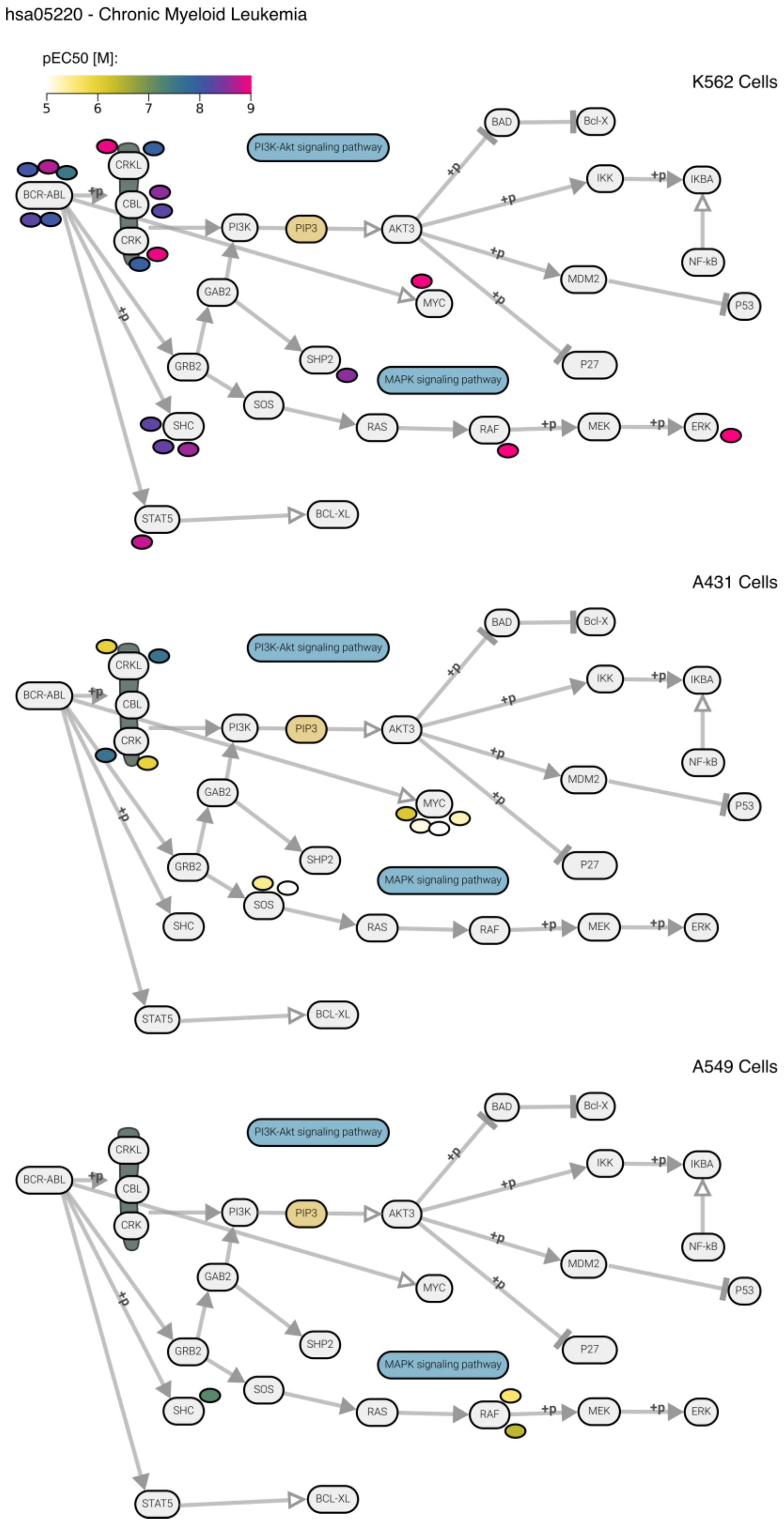


**Fig. S9. Comparison of dose-dependent Dactolisib treatment in three different cell lines.** Edited screenshots from PTMNavigator. Projection of data from Zecha et al. onto the KEGG diagram 'hsa05220 – Chronic Myeloid Leukemia'. Edges labeled with '+p' indicate a phosphorylation of the target protein. *Top:* K562 lymphoblast cells. *Middle:* A431 epidermoid carcinoma epithelial cells. *Bottom:* A549 lung adenocarcinoma cells.

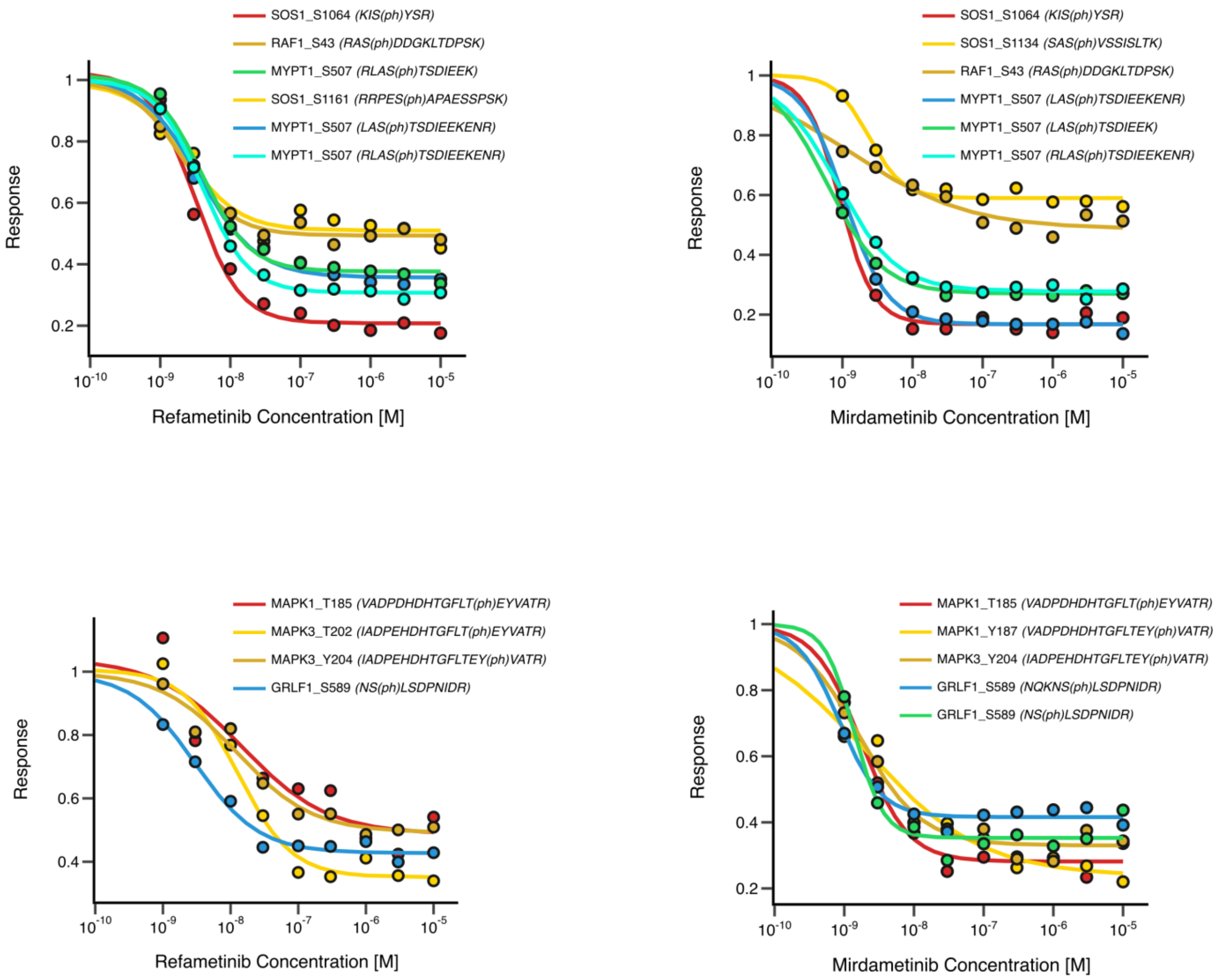


**Fig. S10. decryptM curves from the Refametinib and Mirdametinib datasets.**
